# Virulent but not temperate bacteriophages display hallmarks of rapid translation initiation

**DOI:** 10.1101/2021.04.21.440840

**Authors:** Adam J. Hockenberry, David C. Weaver, Claus O. Wilke

**Affiliations:** Department of Integrative Biology, The University of Texas at Austin

## Abstract

Bacteriophages rely almost exclusively on host-cell machinery to produce their proteins, and their mRNAs must therefore compete with host mRNAs for valuable translational resources. In many bacterial species, highly translated mRNAs are characterized by the presence of a Shine-Dalgarno sequence motif upstream of the start codon and weak secondary structure within the start codon region. However, the general constraints and principles underlying the translation of phage mRNAs are largely unknown. Here, we show that phage mRNAs are highly enriched in strong Shine-Dalgarno sequences and have comparatively weaker secondary structures in the start codon region than host-cell mRNAs. Phage mRNAs appear statistically similar to the most highly expressed host genes in *E. coli* according to both features, strongly suggesting that they initiate translation at particularly high rates. Interestingly, we find that these observations are driven largely by virulent phages and that temperate phages encode mRNAs with similar start codon features to their host genes. These findings apply broadly across a wide-diversity of host-species and phage genomes. Further study of phage translational regulation—with a particular emphasis on virulent phages—may provide new strategies for engineering phage genomes and recombinant expression systems more generally.

## Introduction

Protein translation consumes a large amount of cellular resources, and the genomes of microbial species show strong evidence of selection for rapid and efficient translation [1–8]. The statistical analysis of genome sequence features has provided important insights into many of the basic mechanisms of gene expression, particularly by contrasting sequence patterns within genes that have high- and low-expression demands [9–14]. Over-representation of a purine-rich motif upstream of bacterial start codons, for instance, ultimately lead to the discovery of the Shine-Dalgarno sequence mechanism—a direct binding interaction between the 30S ribosomal sub-unit and mRNA that governs start codon recognition and facilitates translation initiation [15–21]. Collectively, gene sequence analyses have facilitated advances in recombinant protein engineering by deciphering the sequence rules for optimal expression, which have subsequently been validated, refined, and expanded upon in numerous experimental and evolutionary studies [22–32]. However, the amount of available information contained within individual bacterial genomes is limited both in terms of the overall number of proteins and the sequence-level diversity that is present across closely related strains.

Bacteriophage genomes, by contrast, contain a large pool of untapped sequence diversity that metagenomic techniques have only recently begun to characterize in depth [33–39]. Phage mRNAs must be expressed in host cells—in most cases, entirely by existing host cell machinery—and the statistical patterns and constraints that are encoded in these sequences may thus help to further elucidate host-cell transcriptional and translational constraints and mechanisms [40, 41]. While several model phage species are well-characterized [42–46], there have been comparatively few investigations into larger-scale statistical patterns present within phage genomes [47–53]. Deciphering the coding sequence rules governing phage genome evolution is additionally critical for engineering phages as well as for determining the utility of this knowledge for host-cell engineering applications [54, 55].

Here, we analyze thousands of complete, high-quality phage genomes from 33 different bacterial hosts to determine whether the constraints shaping translation-initiation regions in phage mRNAs differ from host-cell genomes. We first present an in depth analysis of *E. coli*-infecting phages and find that phage genomes are predicted to have translation-initiation rates that are on-par with host-cell genes encoding only the mostly highly abundant proteins. Next, we show how translation initiation-related features co-vary and associate strongly with phage lifestyle. Finally, we extend our findings across a broad phylogenetic range of host species, suggesting that phage mRNAs—and virulent phages in particular—are subject to strong evolutionary pressure to ensure rapid translation initiation.

## Results

### Phage mRNAs are predicted to have rapid translation-initiation rates

We focused our study on two sequence features that are robustly linked to translation-initiation rates across a wide-range of bacterial species. For a given gene, we first measured the strength of anti-Shine-Dalgarno (aSD) sequence binding interaction (5’-CCUCCU-3’) by selecting the strongest binding hexamer sequence within a narrow window upstream of the annotated start codon (Fig. 1A). Stronger aSD sequence binding (more negative △*G*) is associated with start codon recognition and rapid translation initiation [56]. Next, we calculated the structural accessibility of the start codon by predicting the stability of the secondary structure for a 90 nucleotide fragment surrounding each start codon (30 bases upstream and 60 bases downstream, Fig. 1B). Weaker secondary structure within this region (more positive Δ*G*) is robustly associated with higher translation-initiation rates [57].

**Figure 1.**
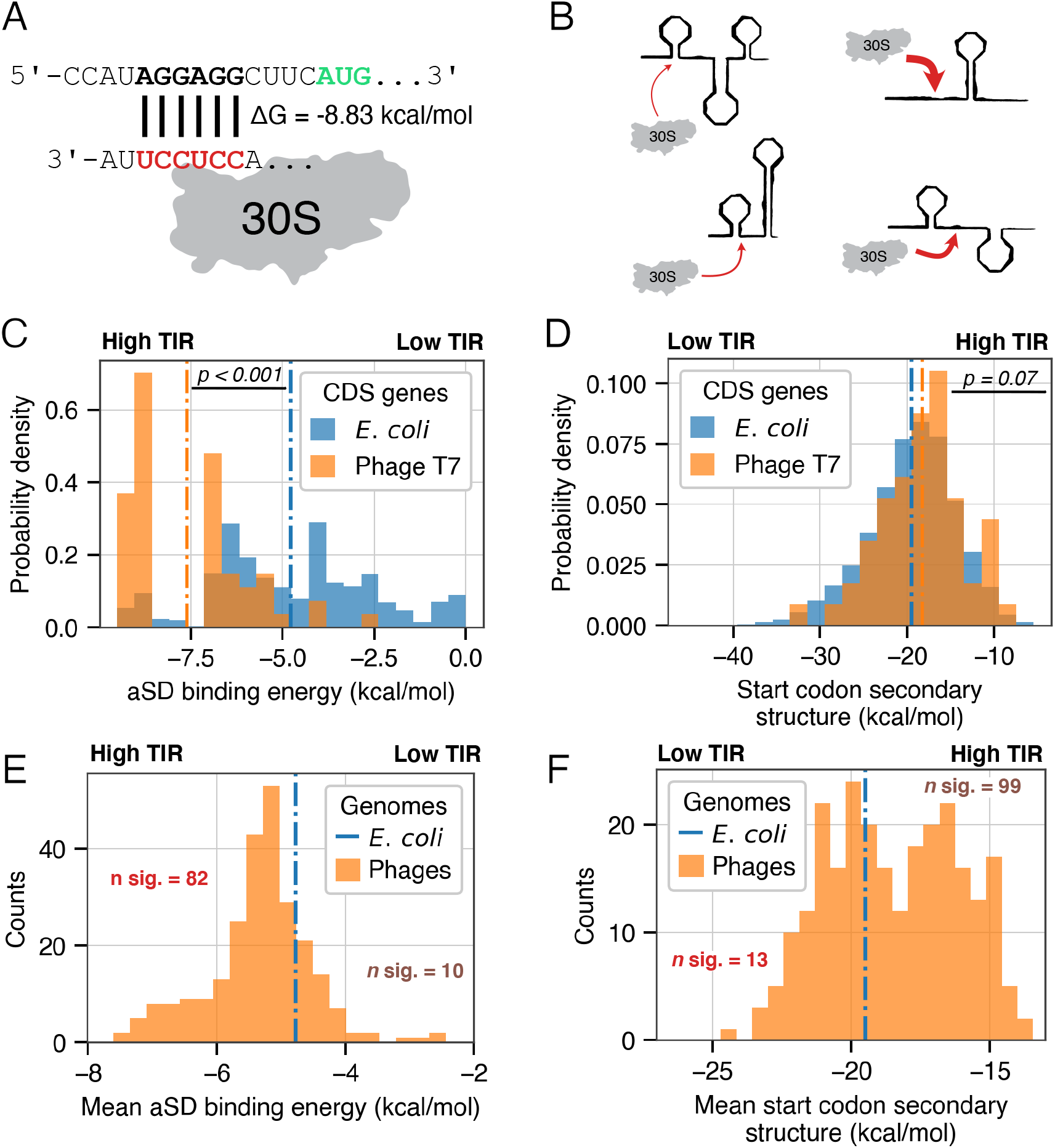
Translation-initiation metrics in phage and host genomes. (A) Illustration of the aSD binding strength, where stronger ribosomal binding is associated with a higher predicted translation-initiation rate (TIR). (B) Illustration of mRNA secondary structure around the start codon, where weak structure is associated with higher predicted TIRs. (C) Histogram of aSD sequence binding strengths upstream of phage T7 and *E. coli* start codons (dashed lines indicate group means). (D) As in (C), showing mRNA secondary structure strength surrounding the start codon. (E) Histogram of mean aSD sequence binding energy across 254 phage genomes. ‘n. sig’ denotes the number of phage genomes with significant differences compared to *E. coli* (dashed line). (F) As in (E), comparing mean mRNA secondary structure around the start codon.

We measured aSD sequence binding and secondary structure strengths for all protein coding genes in phage T7—a well-studied, model phage–and its bacterial host, *E. coli*. We observed a clear distinction whereby the aSD sequence binding strengths of T7 genes are narrower and highly skewed towards stronger binding relative to *E. coli* genes (Fig. 1C, mean of −7.61 vs −4.77 kcal/mol, Welch’s t-test *p* < 0.001). The distribution of secondary structure strengths within the start codon region are slightly (but insignificantly) shifted towards weaker secondary structure, which is also associated with higher translation-initiation rates (Fig. 1D, mean of −18.3 vs −19.5 kcal/mol, *p* = 0.07). Given that aSD sequence binding strengths are substantially stronger and that the strength of mRNA secondary structures are statistically similar in phage T7 relative to *E. coli*, this analysis shows that overall translation-initiation rates (TIR) are predicted to be higher for phage T7 mRNAs relative to host mRNAs.

To determine the generality of these findings, we extended this analysis to a set of 254 diverse *E. coli*-infecting phage genomes (see Materials and Methods). Similar to phage T7, we found that 202 of these 254 phage genomes had a stronger mean aSD sequence binding strength when compared to the *E. coli* genome and 82 of these comparisons were significant (Fig. 1E, Welch’s t-test with FDR-correction, *p* < 0.01). By contrast, only 10 of the phage genomes displayed significantly weaker aSD binding strengths when compared with *E. coli* (according to the same significance criteria).

When we considered mRNA secondary structure surrounding the start codon, we found that 152 out of 254 tested phage genomes had *weaker* mRNA secondary structure in the start codon region compared with *E. coli* genes and 99 of these comparisons were statistically significant (Welch’s t-test with FDR-correction, *p* < 0.01, Fig. 1F). As with aSD sequence binding, this result indicates that phage mRNAs are likely to initiate translation more rapidly than host-cell mRNAs. Only 13 phage genomes had significantly stronger predicted mRNA secondary structures within the start codon region compared to host genome mRNAs (and thus lower predicted translation-initiation rates).

One possible explanation for why phage mRNAs appear to have higher translation-initiation rates than hosts is that phage genomes are highly compact and may be devoid of genes that are rarely expressed or under weak selective pressures—as may be the case for much of the > 4, 000 protein coding genes within the *E. coli* genome. We therefore repeated the above analyses, comparing phage genomes to subsets of the *E. coli* genome with progressively more stringent cutoffs in terms of their average protein abundances (see Materials and Methods). As expected, we found that *E. coli* mRNAs that encode highly abundant proteins have both substantially stronger aSD sequence binding strengths and weaker mRNA secondary structure in the start codon regions. Achieving parity in terms of roughly balancing the number of phage genomes with significantly stronger (and weaker) aSD sequence binding strengths required considering only the top 10–25% *E. coli* genes with the highest protein abundances (Supplementary Fig. S1A). The results when assessing mRNA secondary structure in the start codon region are similar: overall the translation-initiation region of phage mRNAs appear to be under strong selection that is on par with only the most highly expressed host genes (Supplementary Fig. S1B). We additionally considered essential and non-essential host gene categories separately under the expectation that phage genes might more closely resemble essential host-cell genes. However, we found that phage translation-initiation regions are significantly distinct from both essential and non-essential gene subsets (Supplementary Fig. S1A,B).

### Sequence features in translation-initiation regions differ between virulent and temperate phages

Our findings thus far show that phage mRNAs are predicted to bind strongly to ribosomes and thus are predicted to have increased translation-initiation rates relative to host mRNAs. Additionally, phage mRNAs also appear to have generally weaker secondary structures surrounding the start codon, a feature that is itself associated with rapid recruitment of ribo-somes and high translation-initiation rates. However, we have investigated these features in isolation and it is unclear whether these two sequence features represent competing strategies that different phages use to ensure rapid translation initiation or whether these features co-occur within the same genomes.

To investigate the possible differences between phage and host mRNAs along several dimensions, we used a logistic regression framework that attempts to predict the genome of origin (host or phage) based on knowledge of both the aSD sequence binding strength and the strength of mRNA secondary structure surrounding the start codon for each phage-host pairing. Using only a single predictor variable, this procedure is equivalent to a Student’s t-test but the flexible nature of the model allows us to analyze both variables simultaneously and to further account for potentially confounding variables. The reported effect size is simply the model coefficient for each variable (a standardized conditional log-odds ratio), the magnitude of which can be directly compared for various predictor variables to assess their overall contribution. We decided to control for two potentially confounding variables: coding sequence GC content and codon usage biases. The GC content of coding sequences can directly influence the strength of mRNA secondary structure (and this feature may vary between phage and host genomes). Additionally, codon usage biases are a general indicator of translational selection—particularly on translation elongation. We found that GC content and codon usage biases do indeed vary between host and phages when these variables are considered in isolation (Supplementary Fig. S2). Our logistic regression model thus contains two features of direct interest related to translation initiation and two features that we treat as potentially confounding variables.

Using the same dataset of 254 *E. coli* infecting phages, we found that strong aSD sequence binding strength and weak mRNA secondary structure co-occur in a large subset of phages (upper-left quadrant, Fig. 2A). The number of significant phages indicated in the figure (37 in the upper-left quadrant) refers to to phages where *both* variables are significantly different from the host (after separately applying a FDR-correction to each variable, *p* < 0.01). However, numerous phage genomes reside in the lower-left quadrant where they display slightly stronger aSD binding strengths than the host genome but also slightly stronger mRNA secondary structure in the start codon region. Only two of these genomes are significant for both quantities, however. Finally, a comparatively small number of genomes occupy quadrants on the right side of this scatter plot (displaying weaker aSD sequence binding strengths) and only a single phage genome is statistically different from the host for both features—in this case, having significantly weaker aSD sequence binding and stronger mRNA secondary structure in the start codon region.

**Figure 2.**
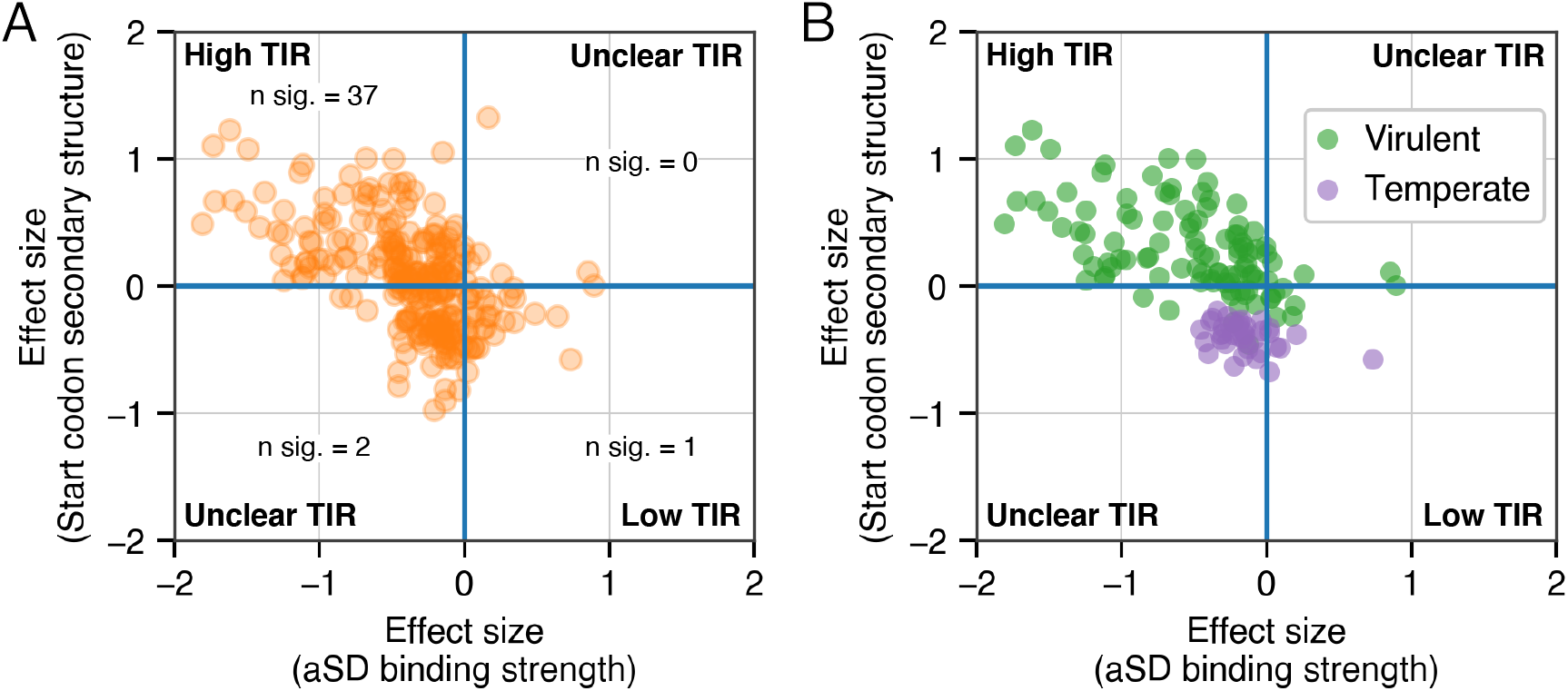
Multi-variable modeling of translation-initiation differences between phage and host genomes. (A) Model coefficients (standardized conditional log-odds ratio) from logistic regression comparing sequence features in translation-initiation regions between individual phage and *E. coli* genomes. Coefficients for aSD binding energies and mRNA secondary structure in the start codon region are depicted on the x- and y-axes, respectively, while coefficients for confounding variables (GC content and codon usage biases) are not shown. (B) As in (A), with phages colored according to their predicted lifestyle for the subset of phages with high-confidence lifestyle predictions (≥ 0.95 probability of correct assignment using BACPHLIP).

All of the results that we have presented thus far treat phage genomes uniformly. However, this approach may obscure important differences in how natural selection operates on the translation-initiation regions of different phages. We thus predicted the lifestyle of each phage genome in the dataset using BACPHLIP [58] and noticed a striking difference between phages that are confidently predicted (≥ 0.95% probability) to be either temperate or virulent (*n*=143, of which 39 are temperate and 104 are virulent, Fig. 2B). While both categories of phages have strong aSD sequence binding energies (most data points remain to the left of zero in Fig. 2B), the virulent phages reside in the upper-left quadrant and almost exclusively have high predicted translation-initiation rates (Fig. 3B). By contrast, the vast majority of temperate phages reside in the lower-left quadrant where they display signatures of strong aSD sequence binding strengths but also stronger mRNA secondary structures as compared to the host. Repeating the full analysis from Fig. 2 without including the coding sequence GC percent and iCUB covariates reveals similar findings, with virulent phages preferentially residing in the top left quadrant, which is indicative of storng predicted translation initiation rates (Supplementary Fig. S3).

**Figure 3.**
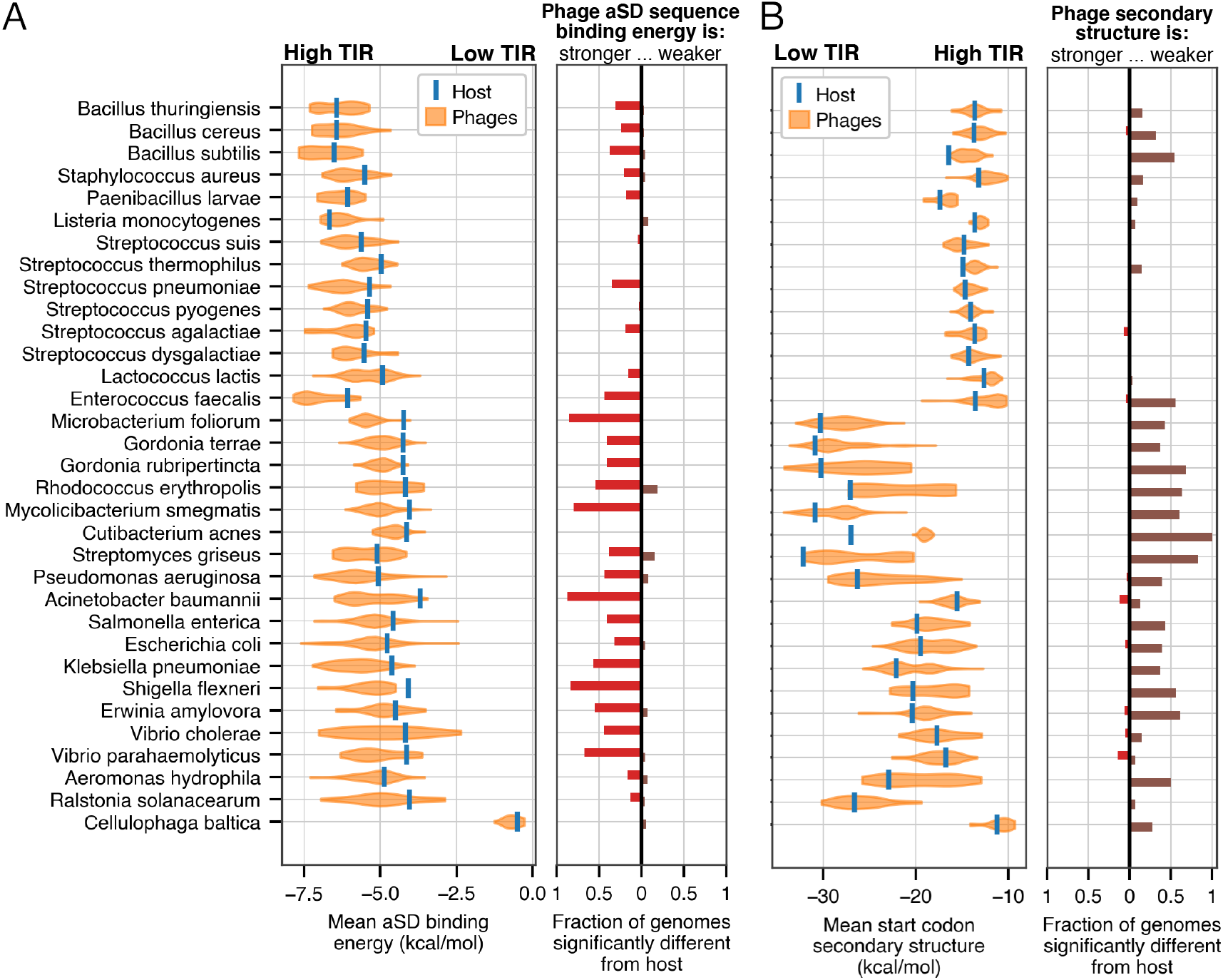
Phage genomes show evidence of high translation-initiation rates across a range of taxa. (A) The distribution of mean aSD binding strengths across all phages that infect an individual host is depicted as a violin plot with the host genome average shown as a blue vertical line. The fraction of phage genomes that are significantly different for each host are depicted in the vertical bar plot on the right, with red bars extending to the left of 0 indicating genomes with significantly stronger aSD binding strengths and brown bars to the right indicating significantly weaker aSD binding strengths (significance was defined via Welch’s t-test with FDR-correction, *p* < 0.01)(B) Similar to (A), results for mRNA secondary structure in the start codon region. For both panels, results for *E. coli* are identical to those depicted in Fig. 1E, F.

We further confirmed this stark separation between temperate and virulent phage genomes by looking at each of the features independently within the temperate and virulent phage genome sets (Supplementary Fig. S4). The clearest and most significant differences emerged when considering mRNA secondary structure around the start codon and codon usage biases. In both cases, virulent phages are predicted to have stronger translation initiation rates and translation efficiency when compared with temperate phages (Welch’s t-test, *p* < 0.001). The strength of aSD sequence binding also differs slightly (*p* = 0.011) between these two sets of phages: of the 143 genomes with confident lifestyle predictions, the top 35 phages with the strongest mean aSD sequence binding energies are all virulent phages. Finally, coding sequence GC content also varied, virulent phages displaying a generally lower coding sequence GC percent (*p* < 0.001) but the link between this feature and translation efficiency is less well studied.

Finally, to better understand how these two translation initiation-related features contribute to translation efficiency—a metric that is roughly akin to the amount of protein produced from a given level of mRNA over a given time period and is often calculated using ribosome profiling. We built multi-variate regression models to predict gene-specific translation efficiencies based off of two previously published, empirically determined values [59, 60]. Both translation initiation features were highly significant, and the resulting models had Adjusted-*R*^2^ values between 0.11-0.17 (Supplementary Fig. S5). We applied the best fitting model to predict translation efficiency values for phage genomes, and observed that a large number had significantly greater predicted mean translation efficiency when compared with the predictions for the entire *E. coli* genome. Confirming our lifestyle-based findings, this effect was driven almost entirely by virulent phage genomes—many of which had significantly greater predicted translation efficiency whereas no temperate phage genomes showed this effect (Supplementary Fig. S5).

### High translation-initiation rates are a general feature of phage genomes

We have thus far presented an in depth analysis of *E. coli* infecting phage genomes, but the generality of these findings to other host-cell bacterial species is unclear. To determine if phage mRNAs display evidence of rapid translation initiation in general, and whether this effect is particularly pronounced in virulent phages, we repeated our analyses from Fig. 1E, F on 32 additional taxonomically diverse bacterial host organisms for which we have at least 10 (dereplicated) complete phage genomes for comparison.

For nearly all host species, we found that a substantial fraction of phage genomes had significantly different aSD sequence binding strengths and mRNA secondary structure strengths compared to their hosts (Fig. 3, the fraction of significant genomes depicted in the bar chart is calculated via Welch’s t-test with FDR-correction, *p* < 0.01). In nearly all cases, the effect was heavily biased in the direction of increased predicted translation-initiation rates for phage mRNAs: stronger aSD sequence binding strengths and weaker mRNA secondary structure in the start codon region. There were, however, a few scattered host species for whom there appeared to be no significant difference for one or both of the translation-initiation features. In *Cellulophaga baltica*, for instance, we note that it has previously been shown that many members of the *Bacteroidetes* phylum do not use the aSD sequence binding mechanism and it is thus unsurprising to see that phages infecting this species are also devoid of this feature [6, 61, 62]. Additionally, we observed generally weaker effects for both translation initation-related features across members of the *Firmicutes* phylum (excepting the *Bacillus* genus) but note that the translation-initiation regions in many of these host-cell genomes have particularly strong features compared with species like *E. coli*.

We additionally wished to assess differences between temperate and virulent phages for this diverse set of species. However, most phages with known, annotated lifestyles come from a biased set of host species and this bias also affects the accuracy of phage lifestyle *prediction* for different host-cell species. We predicted the lifestyle of all phages infecting the host-cell species depicted in Fig. 3 and (as in Fig 2) required a 95% probability of correct lifestyle assignment. This procedure yielded 9 host species for which we could confidently identify a minimum of 5 temperate and 5 virulent phage genomes. For these hosts, we split the phage genomes into temperate and virulent categories and repeated the analysis from Fig. 3 while considering temperate and virulent phages independently.

Similar to our findings across *E. coli* infecting phages, a substantial number of both virulent and temperate phages displayed stronger aSD sequence binding strengths relative to host-cell genomes and we observed comparatively minor differences in the magnitude of this feature across lifestyle classes Fig. 4A. Also confirming our results seen in *E. coli*, we observed very few cases where temperate phages were predicted to have significantly different mRNA secondary structure in the start codon region in either direction. By contrast, large numbers of virulent phages have weaker mRNA secondary structure in the start codon region Fig. 4B. Virulent phages, across nearly all of the 9 species studied, have strong sequence signatures that are indicative of rapid translation initiation. As with *E. coli*, the picture is more nuanced for temperate phages, which tend to have strong aSD sequence binding strengths (Fig. 4, left) but little or no significant difference in mRNA secondary structure strengths in the start codon region compared to host genes (Fig. 4, left).

**Figure 4.**
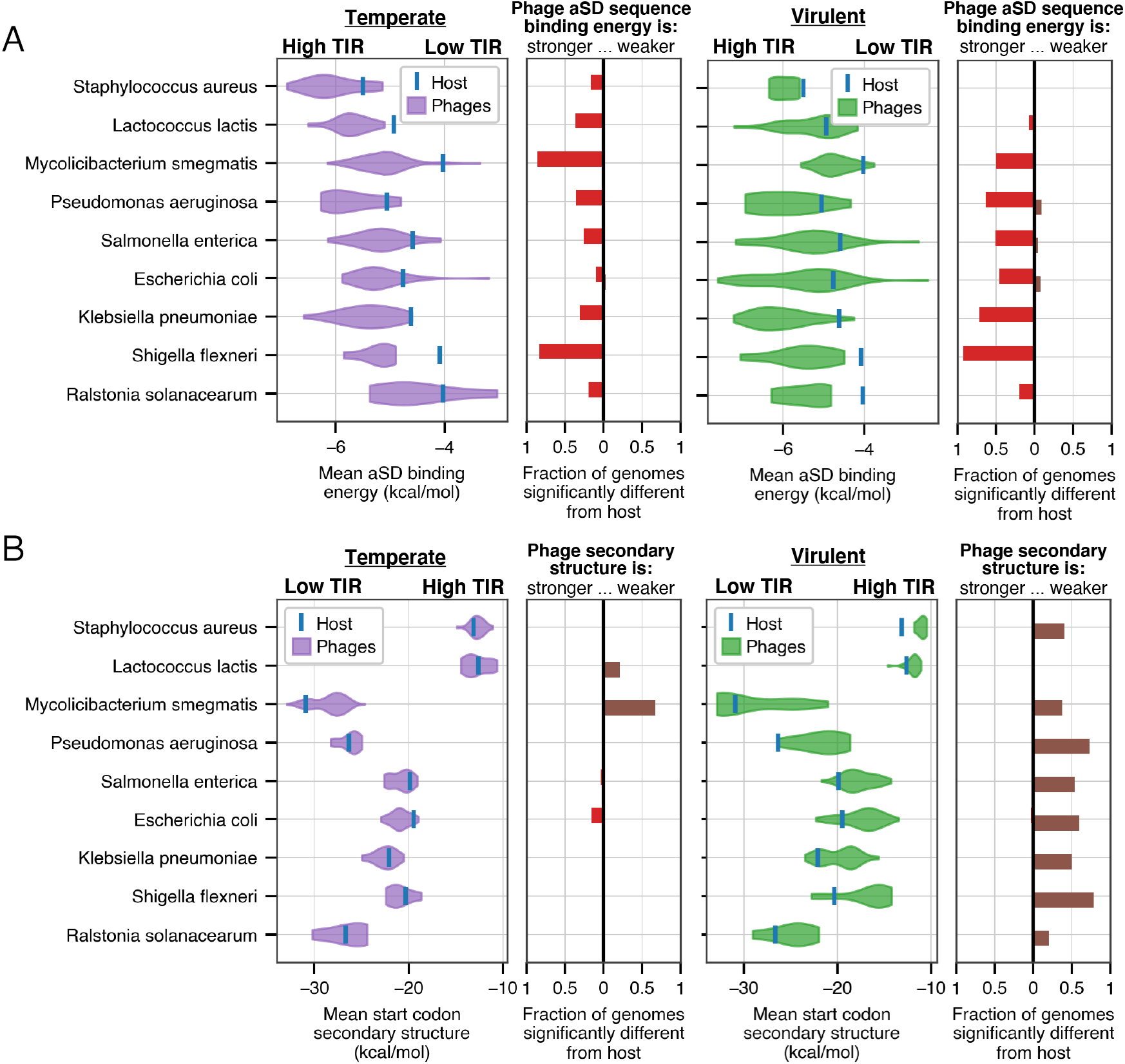
Phage lifestyle and translation initiation sequence feature variation across diverse bacterial species. (A) Comparing mean host-genome aSD sequence binding energies independently with values from temperate (left, purple) and virulent (right, green) phage genomes. Most virulent and temperate phage genomes across all host-cell species display strong aSD sequence binding. (B) Comparing mean host-genome secondary structure energies in the start codon region with values from temperate (left, purple) and virulent (right, green) phage genomes. Virulent phages, but not temperate phages, have weaker mRNA secondary structure in the start codon region across a range of host-species.

## Discussion

Translation initiation region sequence preferences provide bacteria with a means to differentiate start codons from background sequences and to modulate the protein production rate for individual mRNAs. By analyzing the sequence patterns contained within thousands of complete bacteriophage genomes, we have shown these sequence preferences are particularly strong in phages and indicative of rapid translation-initiation rates that are on par with only the most highly expressed host-cell genes. Most strikingly, we find that virulent phage genomes differ from temperate phage genomes in this regard: mRNAs from both types of phages appear to be enriched in strong anti-Shine-Dalgarno sequence binding sites, but only virulent phages couple this ribosomal capture mechanism with particularly weak mRNA secondary structure that helps to facilitate ribosomal recruitment.

Within temperate phage genomes, the net result of partially contradictory observations—stronger aSD sequence binding and slightly stronger mRNA secondary structure relative to host genes—is unclear. To date, there have been a limited number of genome-wide transcriptomic and proteomic studies specifically targeting phage infections [42–46]. As techniques such as RNA-seq and ribosome profiling become more widespread and applied to diverse phage and host species, we expect that the details of ribosomal competition during phage infection will become more clear. However, based on regression models that were fit to empirically determined *E. coli* translation efficiency values, it appears that temperate phages are likely to have slightly lower translation efficiency (Supplementary Fig. S5). By contrast, virulent phage genomes appear to have unequivocally increased rates of translation initiation according to all of our best knowledge about this detailed molecular process. Our study thus joins a growing body of literature highlighting important differences in evolutionary processes between temperate and virulent phage genomes [63–68].

While we are unable to say precisely *why* temperate and virulent phages should differ so starkly in regards to their translational initiation regions, we speculate that the life-history of these different virus types results in unique translational pressures. Temperate phage genomes, for instance, may accumulate numerous point mutations that are likely to be under weak selection pressure during extended periods of dormancy. It is possible that excision of temperate phages from the host-cell genome (and subsequent entry into the lytic cycle) is preceded by other steps that limit competition between phage and host-cell mRNAs. By contrast, virulent phage genomes must rapidly produce protein in a race against host-cell detection mechanisms in order to reproduce at every generation. While not the focus of our study, we observed that codon usage biases also differ significantly between temperate and virulent phages, with temperate phages having generally weaker biases than we observed for virulent phage genomes. This again points in the direction of stronger overall levels of translational selection on virulent phages. However, the precise consequences of these different evolutionary modes as it relates to optimizing translation initiation remains an intriguing area of further theoretical and empirical study.

Prior work has shown that viral mRNAs (including phage) tend to have weak mRNA secondary structure surrounding the start codon, but limits in the number of available phage genomes made it difficult to explicitly study phage-host pairings [69]. Other studies have looked at differences in codon usage biases and higher-order sequence effects within bacterio-phage genomes and have generally found strong signatures of translational selection in phage genomes [47–53]. Our findings thus build on existing research and suggest that a distinct an important comparison is between host and phage mRNAs, which are likely to be in direct competition for the same pool of ribosomes and translational resources during periods of active infection. The growing abundance of complete, host-linked phage genomes presents exciting opportunities for further study in this area.

A potential limitation of our work is that phylogenetic relatedness presents numerous statistical challenges. Two randomly chosen phages may share either close or distant sequence identity and common ancestry [70]. While phylogenetic comparative methods can be applied to correct for non-independence, these methods require an underlying phylogenetic tree that can be particularly challenging to assemble for phage genomes [39, 71, 72]. In lieu of this approach, we have opted for a simpler method that should partially mitigate phylogenetic biases: clustering genomes and selecting single representative genomes per cluster to ensure that the data points are more independent than they might otherwise be. Additionally, while we present multi-species comparisons between various host-cell species (Figs. 3 & 4), we again have not accounted for any confounding effects that could arise due to phylogenetic relatedness between host-cell species. However, we do not perform any explicit statistical analyses on the entire set of data, and rather include these figures to graphically show that our findings appear to apply across numerous species.

Analysis of protein coding sequences, as well as up- and down-stream regulatory sequences, has enhanced our understanding of several transcriptional and translational mechanisms, including the role of codon usage biases and higher-order sequence features in the rate of protein production [73–78]. In a sense, phages are the original synthetic biologists—exploiting host cells to produce a set of macromolecules that decrease host fitness. The precise details of how individual phages manipulate their hosts is highly varied, but higher-order genome features such as we have studied here may provide important insight into translational regulation, which may further enhance our ability to engineer both phage and host-cell genomes.

## Materials and Methods

### Virus data compilation, genome annotation, and host prediction

The core of our study relies on access to numerous and distinct bacteriophage genomes with consistent annotations and trustworthy predictions of primary host species. On-going research in each of these areas is continuing to expand the availability of phage genomes that are suitable for genome-linked analyses such as we performed here [39, 79–83]. To maintain consistency and to ensure access to the highest-quality genomes, we used the NCBI Virus genome database (last accessed: November 2020) and selected only “complete” genomes to include in our study. An advantage of this choice is that we additionally relied on previously existing genome annotations for all phages included in this study. Only a small subset of phage genomes have host annotations, so we limited our selection of host-cell species to those with at least 50 annotated phage genomes and 10 annotated phage genomes after dereplication (see below).

To ensure that the selected phage genomes were not severely biased by a small number of over-represented and closely related genomes, we used FastANI to measure the average nucleotide identity (ANI) between all phages that infect a single host, requiring that the alignment span a minimum of 80% the length of the shortest genome [84]. We constructed a unique all-to-all distance matrix, and used the cd-hit-est greedy algorithm to cluster phages at 95% sequence identity [85]. From each cluster, we subsequently selected the longest Ref-Seq genome as a cluster representative. In the event that there were no RefSeq genomes in a cluster, we simply selected the longest genome. Finally, we removed poorly annotated phage genomes from the analysis, which we conservatively defined as those with an annotated coding sequence density <50%. For *E. coli*, our processing began with a set of 1,473 genomes and our final dataset encompassed 254 genomes, which were analyzed throughout this manuscript.

### Definitions of sequence features related to translation initiation

The Shine-Dalgarno sequence is a collection of related mRNA sequence motifs that are defined by their ability to bind strongly to the highly conserved anti-Shine-Dalgarno sequence— located on the 30S subunit of the bacterial ribosome. Here, we define the aSD binding strength of a given coding sequence as the strongest possible binding affinity between the anti-SD sequence (defined via the core sequence of 5’-CCUCCU-3’) and hexamer sequences upstream of the start codon, restricting the upstream range to have a minimum gap of 4 nucleotides and a maximum gap of 10 nucleotides between the 3’ most base in the hexamer and the first nucleotide of the start codon. Binding energies were calculated using the “RNAcofold” program, part of the ViennaRNA (v2.4.14) suite [86].

We determined the strength of mRNA secondary structure surrounding all start codons by assessing a 90 nucleotide long sequence fragment (30 nts upstream of the start codon and 60 nts downstream) using the “RNAfold” program from ViennaRNA (v2.4.14) [86]. While we expect this to be a rough approximation of the true secondary structure present in the start codon region for each mRNA, numerous prior studies have showed that similar window sizes roughly capture the relevant feature of mRNA secondary structure strength [22, 56, 57].

More complicated models of translation initiation may include explicit penalties for SD sequence spacing relative to the start codon [23], the identity of the start codon itself [87], kinetics of secondary structure unfolding and refolding [88], penalties for binding too strongly to the aSD sequence [56], as well as a number of other possible features [89–91]. However, it is unknown how transferable these other mechanisms are across diverse organisms given that the vast majority of existing work has been performed in *E. coli*. To keep our model simple, we focused on the two most consistent and frequently cited modifiers of translation-initiation rates. Additionally, we found that a simple multi-variable linear regression model consisting of only aSD binding strength and the strength of secondary structure around the start codon is capable of significantly predicting translation efficiencies derived from two separate genome-wide *E. coli* datasets [59, 60] (Supplementary Fig. S5). In these models, *R*^2^ values ranged from 0.11–0.17 and predictions of both models were highly correlated. Further, in both cases, start codons with weak mRNA secondary structure and strong aSD sequence binding have the highest empirically measured translation efficiency values and both sequence features were highly significant in the regression models (*p* < 0.001). We did not apply these models to predict translation efficiencies for phages or hosts outside of *E. coli*, reasoning that doing so would make a dangerous assumption that the mechanisms of translational regulation remain similar across phylogenetically diverse species. Our current approach does, however, assume that the mechanisms of translation efficiency and translational regulation remain similar between normal *E. coli* growth and during phage infection.

### Other CDS-level feature definitions and inclusion criteria

Codon usage bias is frequently considered an indicator of translational selection due to the potential for synonymous codons to modulate the rate of translation elongation. We thus wanted to ensure that our findings were robust to variation in codon usage bias differences between host and phage genomes, as well as GC content variation (which has a direct impact on mRNA secondary structure strength). Coding sequence GC content was simply calculated for each CDS as the number of G+C nucleotides divided by the total coding sequence length. Codon usage bias was measured using iCUB [92]. There are many possible metrics of codon usage bias, but a benchmark study showed that iCUB outperformed other metrics that rely solely on coding sequence information (as opposed to *a priori* defined reference sets of genes or other external information such as tRNA abundances or a reference set of highly expressed genes). The iCUB metric is similar to the more commonly cited effective number of codons: lower values indicate fewer codons and thus more bias. However, iCUB explicitly controls for GC content variation and produces better estimates of protein abundance across diverse microbial taxa. We observed that both of these features (GC-content and iCUB) differ significantly between phage and host species (Supplementary Fig. S2), providing validation that our results are likely to be more robust and conservative by controlling for these effects.

For all studied coding sequences (either phage or host-derived), we only considered annotated genes whose length was at least 90 nucleotides (30 amino acids) and for which the length was a multiple of three (potentially removing a small number of genes with programmed frame-shifts). Additionally, all of our results for host genomes excluded coding sequences whose “product” feature annotation contained the word “phage”. This filter was performed to ideally exclude prophage-associated coding sequences from being included in the host genome when drawing comparisons.

### Compilation of *E. coli*-specific empirical data

We leveraged several existing genome-scale, empirical data sources to thoroughly contrast *E. coli* infecting phages with various subsets of *E. coli* genes as well as to ensure that our translation-initiation metrics were associated with empirically derived translation efficiency measurements. Specifically, we relied on PAXdb for protein abundance data [93], two separate datasets of gene essentiality [94, 95], and two separate datasets of ribosome-profiling derived translation efficiency measurements (briefly referenced in the preceding section) [59, 60].

For protein abundance data, we simply drew various percentile-based thresholds to consider increasingly stringent sets of “highly-expressed” genes. PAXdb aggregates protein abundance measurements from numerous studies and growth conditions, and while these abundance values are estimates from only a small snap-shot of possible studies and possible growth conditions, these values are well-established (if rough) indicators of overall protein abundance [93]. For gene essentiality, we used a consensus approach where we only considered genes that were considered either “essential” or “non-essential” in two separate datasets (to increase robustness) and did not consider genes with conflicting annotations for this portion of the analysis.

### Phage lifestyle prediction

We used the BACPHLIP software [58] to categorize all phage genomes in our dataset into temperate and virulent lifestyles. BACPHLIP uses genome-sequence input, determines the presence/absence of several hundred lysogeny-associated protein domains, and uses a random forest classifier return a probability of the given phage being either temperate or virulent. Here, we limited our analyses of both *E. coli* infecting phages and phages that infect other species to those with a predicted lifestyle probability ≥ 0.95. Additionally, when assessing differences between temperate and virulent phages, we removed any host species from our analysis that did not have at least 5 phage genomes (after dereplication) in each lifestyle category.

### Statistical tests

Single-variable comparisons were performed in Python using the scipy package implementation of Welch’s T-test. Multiple comparisons were corrected for by using the applying the statsmodels implementation of FDR correction (Benjamini/Hochberg) to the list of *p*—values with alpha set to 0.01. Multivariate analyses were performed using the logit function in statsmodels with zscore normalization to all predictor variables in order to standardize effect sizes. Correction for multiple comparisons in these models was again accomplished by filtering the *p*—values from individual model coefficients through the FDR correction procedure with the alpha value set to 0.01. Thus, we ensured that all of our results were highly significant and robust to artifacts arising from multiple comparisons.

### Host taxonomy assignment

We did not explicitly model any effects across a phylogenetic tree and instead present analyses of multiple species independently in Figs. 3 & 4. We do, however, note that no phage genomes were shared across host species for any part of this analysis, but individual host species nevertheless have their own complicated and non-independent phylogenetic histories. We arranged species taxonomically for easier visual comparison by querying the NCBI taxonomy database and used the ete3 package to roughly order species according to their taxonomic grouping. All statistical results were performed for each species independently.

## Code and data availability

All code and data necessary to re-create the analyses in this manuscript are available at https://github.com/adamhockenberry/phage-translation and https://doi.org/10.5281/zenodo.4708008, respectively.

## Acknowledgements

This work was supported by National Institutes of Health grants R01 GM088344 to C.O.W. and F32 GM130113 to A.J.H. C.O.W. also received support from the Jane and Roland Blumberg Centennial Professorship in Molecular Evolution and the Dwight W. and Blanche Faye Reeder Centennial Fellowship in Systematic and Evolutionary Biology at The University of Texas at Austin. The Texas Advanced Computing Center (TACC) at The University of Texas at Austin provided high-performance computing resources. The authors additionally acknowledge valuable feedback and support from members of the Wilke lab.

## Competing financial interest

The authors have declared that no competing interests exist.

## Supplementary Information

**Supplementary Figure S1.**
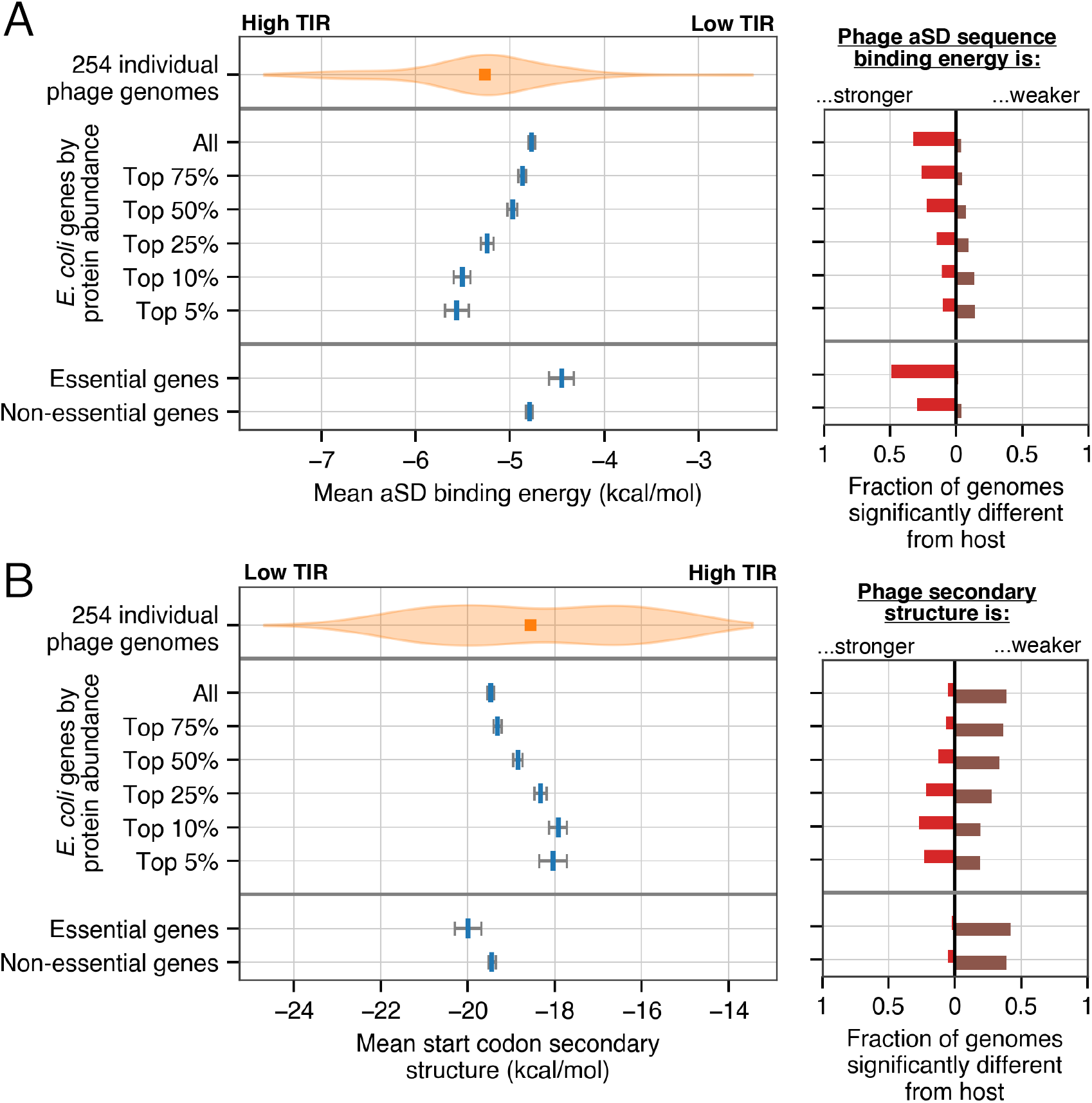
Comparing translation-initiation features in phage genomes to host genome subsets. (A) The distribution of mean aSD binding energy across phage genomes compared with indicated subsets of host genes (error bars depict standard errors for host genome subset means, orange square indicates the median phage genome value). The number of phage genomes that are significantly different from gene categories are shown on the right, with red bars extending to the left of 0 indicating genomes with significantly stronger aSD binding strengths and brown bars to the right indicating significantly weaker aSD binding strengths (significance was defined via Welch’s t-test with FDR-correction, *p* < 0.01). (B) As in (A), depicting mean start codon secondary structure around the start codon.

**Supplementary Figure S2.**
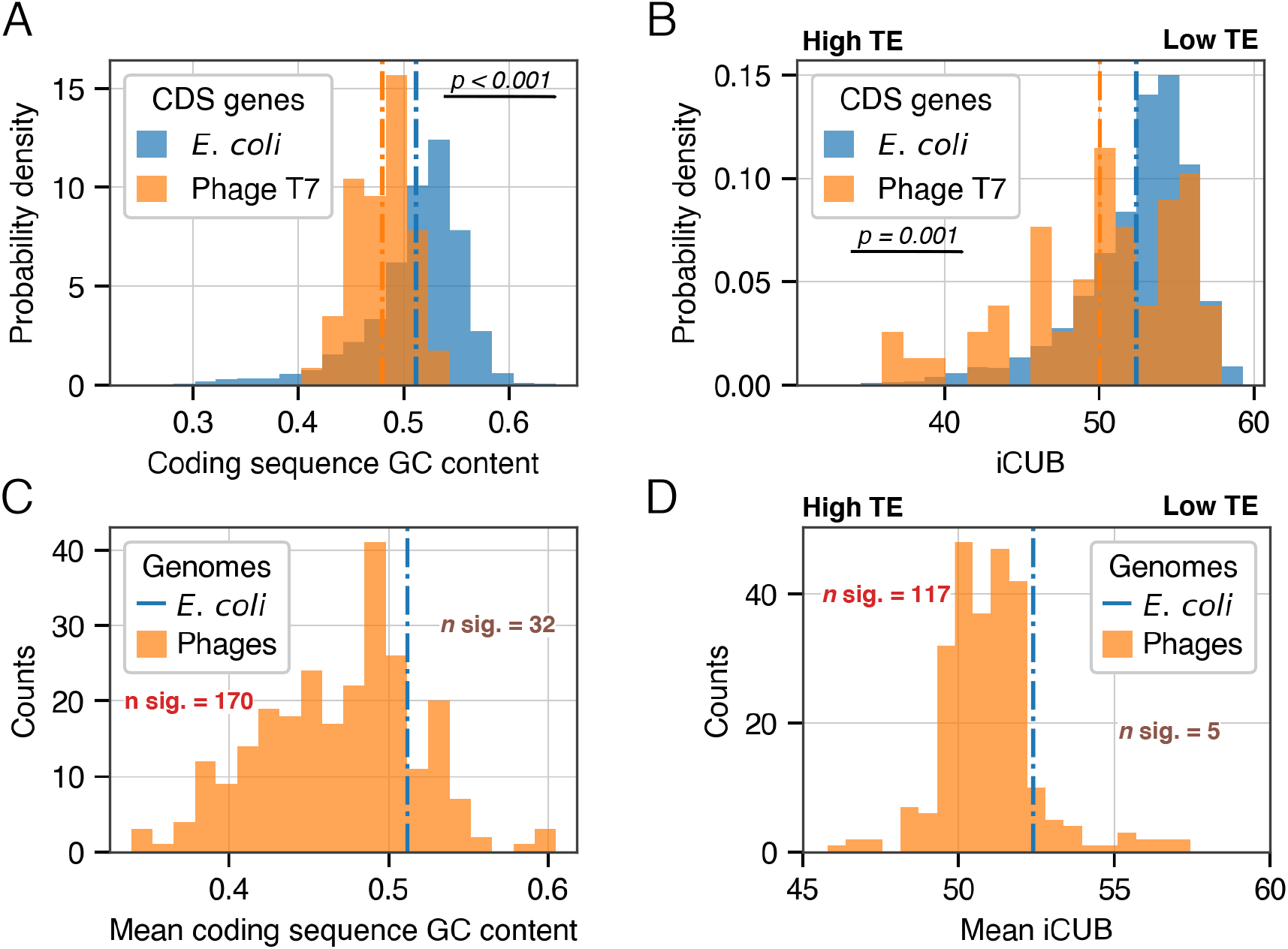
GC content and codon usage bias variation in *E. coli* infecting phage genomes. (A) GC content of all phage T7 and *E. coli* coding sequences. (B) As in (A), showing codon usage bias variation (measured with iCUB). (C) Distribution of mean GC contents across all phage genomes, with the host genome mean depicted as a dashed blue line and the number of significant comparisons on either side annotated as ‘n. sig’. (D) As in (C), showing the mean iCUB values for each phage genome contrasted with the *E. coli*. In panels (B) and (D) the abbreviation TE refers to translation efficiency, indicating that lower iCUB values are associated with stronger translational selection. A similar correspondence GC content and translation efficiency or translational selection is not known.

**Supplementary Figure S3.**
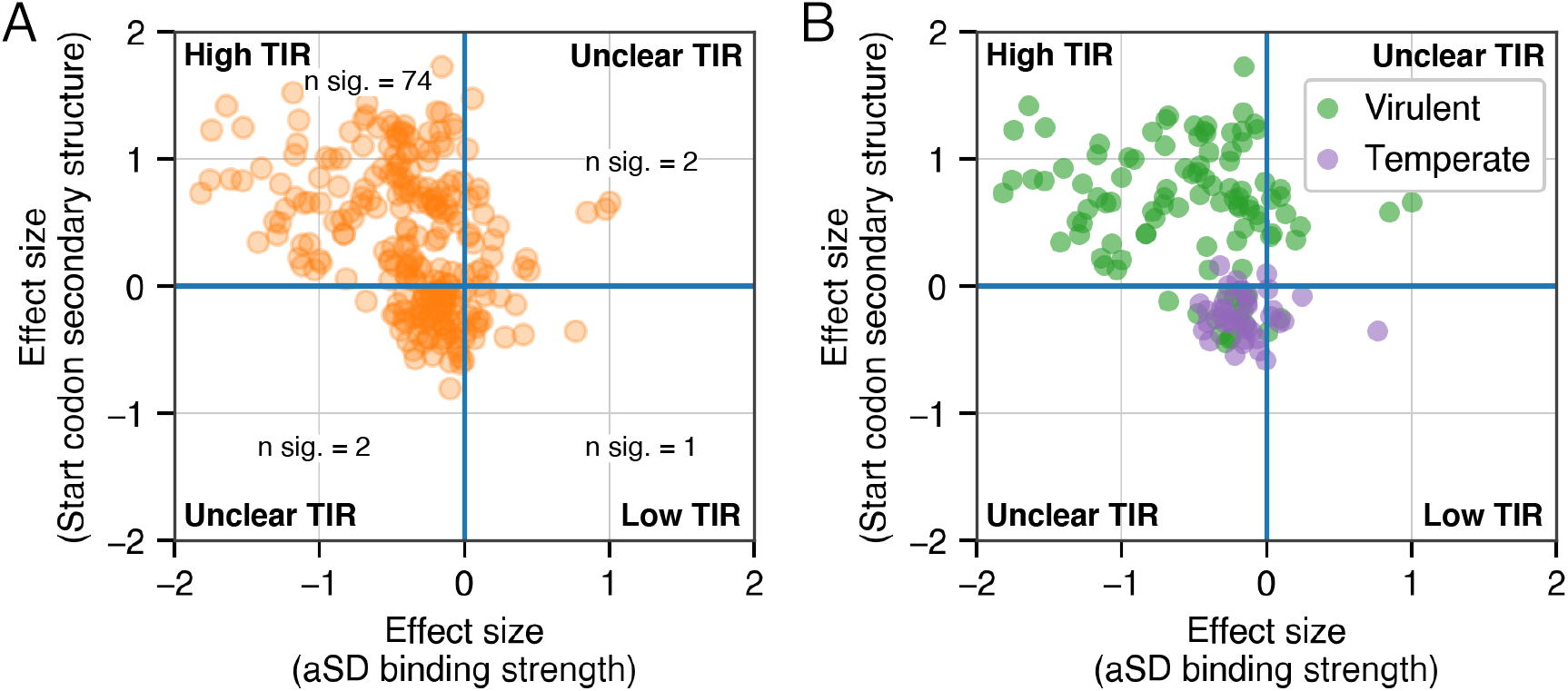
Multi-variable modeling of translation-initiation differences between phage and host genomes without covariates. (A) Model coefficients (standardized conditional log-odds ratio) from logistic regression comparing sequence features in translation-initiation regions between individual phage and *E. coli* genomes. Coefficients for aSD binding energies and mRNA secondary structure in the start codon region are depicted on the x- and y-axes, respectively. (B) As in (A), with phages colored according to their predicted lifestyle for the subset of phages with high-confidence lifestyle predictions (≥ 0.95 probability of correct assignment using BACPHLIP).

**Supplementary Figure S4.**
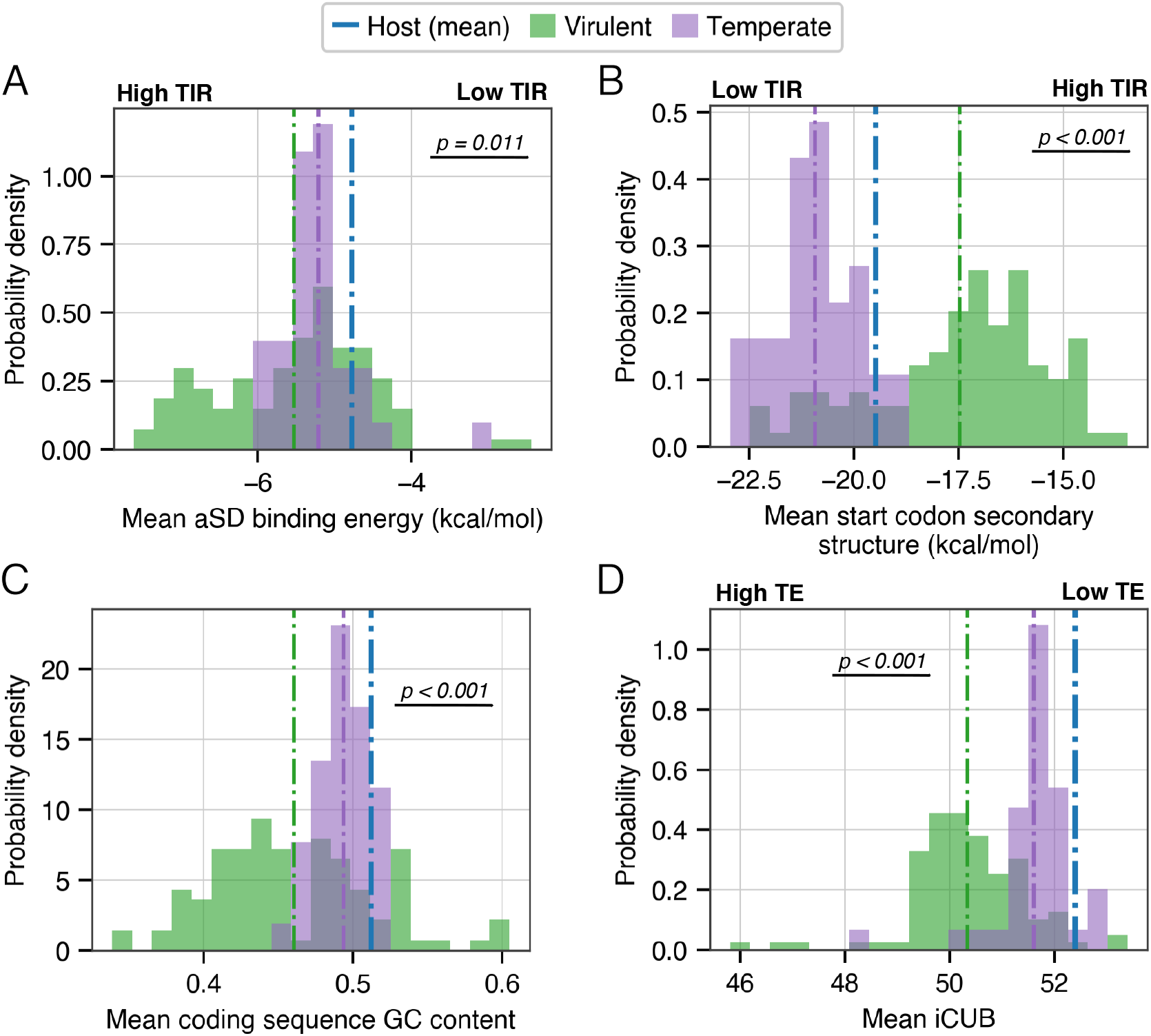
Single variable distributions for 143 E. *coli*-infecting phages with confident lifestyle predictions. (A) Mean aSD sequence binding energy, with virulent phages shown in green and temperate phages in purple. In both cases the dashed colored lines show distribution means, and the dashed blue line indicates the *E. coli* genome-wide mean. Further subplots show: (B) mean start codon region secondary structure (C) mean coding sequence GC percentage and (D) mean iCUB values (akin to the effective number of codons with lower values representing stronger codon usage biases and increased predicted translation initiation rates. In all cases, indicated p-values represent Welch’s t-test comparing temperate and virulent phage distributions (with the host mean shown only as a reference).

**Supplementary Figure S5.**
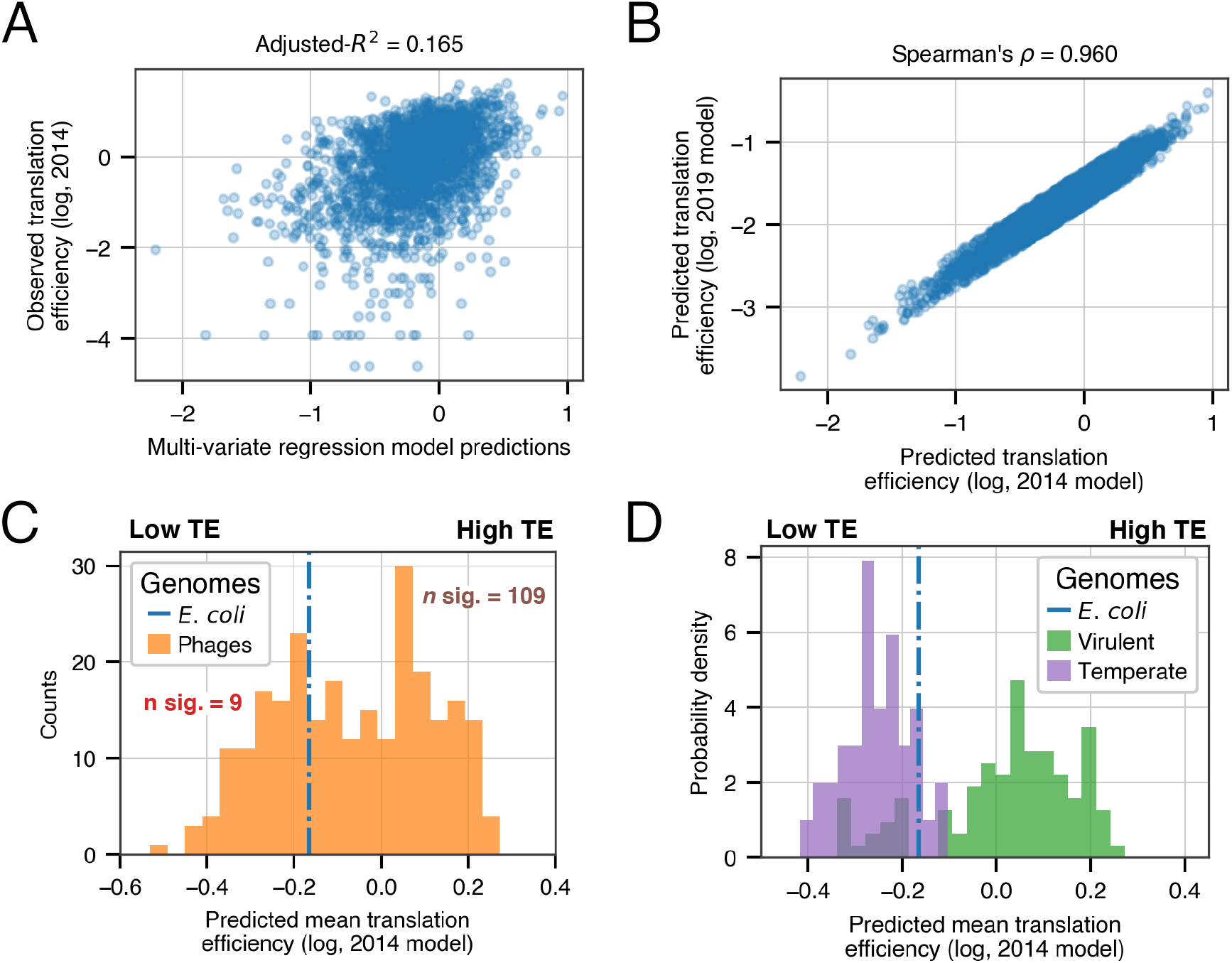
Differences in translation efficiency across phage lifestyles. (A) Scatter plot of predicted vs observed translation efficiency (log-scaled) using translation efficiency data from Li *et al*. (2014) [59] and a multi-variable linear regression model that includes only aSD sequence binding strength and mRNA secondary structure around the start codon. (B) Comparing model predictions across the entire *E. coli* genome when fit to two separate translation efficiency datasets (2014 refers to Li et *al*. (2014) [59] while 2019 refers to Gorochowski et *al*. (2019) [60]. (C) Differences in mean predicted translation efficiency (using the 2014 model) between phage genomes and the host *E. coli* genome. ‘n. sig’ denotes the number of phage genomes that are significantly differenent from *E. coli* in either direction (Welch’s t-test with FDR-correction, *p* < 0.01). (D) As in panel (C), splitting phage genomes according to lifestyle (including only genomes with high-confidence lifestyle predictions: 39 temperate and 104 virulent). Among virulent phage genomes, 73 (0) have significantly higher (and lower) predicted translation efficiency compared to *E. coli*. By contrast, 0 (3) temperate phage genomes have significantly higher (and lower) predicted levels of translation efficiency.

## Notes

https://doi.org/10.5281/zenodo.4708008

### Competing Interest Statement

The authors have declared no competing interest.

